# Investigating the interaction between white matter and brain state on tDCS-induced changes in brain network activity

**DOI:** 10.1101/2020.10.09.332742

**Authors:** Danielle L. Kurtin, Ines R. Violante, Karl Zimmerman, Robert Leech, Adam Hampshire, Maneesh C. Patel, David W. Carmichael, David J. Sharp, Lucia M. Li

## Abstract

**Background:** Transcranial direct current stimulation (tDCS) is a form of noninvasive brain stimulation whose potential as a cognitive therapy is hindered by our limited understanding of how participant and experimental factors influence its effects. Using functional MRI to study brain networks, we have previously shown in healthy controls that the physiological effects of tDCS are strongly influenced by brain state. We have additionally shown, in both healthy and traumatic brain injury (TBI) populations, that the behavioral effects of tDCS are positively correlated with white matter (WM) structure.

**Objectives:** In this study we investigate how these two factors, WM structure and brain state, interact to shape the effect of tDCS on brain network activity.

**Methods:** We applied anodal, cathodal and sham tDCS to the right inferior frontal gyrus (rIFG) of healthy (n=22) and TBI participants (n=34). We used the Choice Reaction Task (CRT) performance to manipulate brain state during tDCS. We acquired simultaneous fMRI to assess activity of cognitive brain networks and used Fractional Anisotropy (FA) as a measure of WM structure.

**Results:** We find that the effects of tDCS on brain network activity in TBI participants are highly dependent on brain state, replicating findings from our previous healthy control study in a separate, patient cohort. We then show that WM structure further modulates the brain-state dependent effects of tDCS on brain network activity. These effects are not unidirectional – in the absence of task with anodal and cathodal tDCS, FA is positively correlated with brain activity in several regions of the default mode network. Conversely, with cathodal tDCS during CRT performance, FA is negatively correlated with brain activity in a salience network region.

**Conclusions:** Our results show that experimental and participant factors interact to have unexpected effects on brain network activity, and that these effects are not fully predictable by studying the factors in isolation.

## Introduction

Transcranial direct current stimulation (tDCS) is a powerful clinical and research tool that can improve cognition in healthy [1] and patient populations [2]. However, variability in the efficacy of tDCS, owing to our insufficient understanding and control of sources of intra and interindividual variability, is a hurdle to its widespread clinical application [3]. Participant factors, such as brain structure, and experimental factors, such as stimulation intensity, all influence tDCS’s effects [3]. Multimodal neuroimaging facilitates the direct study of the physiological effects of stimulation. An increased understanding of how different factors influence the physiological effects of tDCS is invaluable to the rapidly developing field of tDCS for cognitive enhancement [4] [5] [6].

Brain state is a key factor determining the physiological response to tDCS. Previous work from our group and others have found that brain state influences the brain network effects of tDCS in healthy controls [7] [8] [9]. Specifically, our previous tDCS-fMRI work shows that tDCS reinforces the underlying pattern of brain network activity, and that this interacts with stimulation polarity [9].

Work from our group [10] [11] and others [12] have also found that WM structure is a key factor in determining the behavioral response to tDCS. Traumatic brain injury (TBI) characteristically results in traumatic axonal injury due to axonal shearing, where forces tear delicate WM axons [13] [14]. This makes TBI a useful model of reduced white matter connectivity. We have previously found that higher WM structural connectivity, as assessed by fractional anisotropy with Diffusion Weighted Imaging, is related to greater behavioral improvements in a cognitive control task in both healthy and TBI participants [10] [11]. However, there is no previous study investigating the interaction of multiple factors on brain network activity.

In this study, using this previously published group of healthy and TBI participants, we deploy a different stimulation and task fMRI protocol to specifically study how these two factors, that independently influence tDCS’s action, interact to shape the brain network effects of tDCS.

We used the Choice Reaction Task (CRT) to manipulate brain state. The CRT is a well-described task that produces robust patterns of anti-correlated activity in the salience network (SN) and default mode network (DMN) in TBI patients and healthy controls [15]. Additionally, the CRT is a simple cognitive task with which both TBI patients and healthy controls can engage with high (>90.0%) levels of accuracy [15], producing the core pattern of SN activation and DMN deactivation in both healthy control and TBI cohorts. In our previous work, stimulation did not improve behavioral performance in healthy participants [9], enabling changes in brain network activity to be interpreted independently of changes in performance.

We targeted anodal, cathodal, and sham stimulation to the rIFG during execution of the CRT and “rest” blocks. The right inferior frontal gyrus (rIFG) is a control node for DMN/SN dynamics[16] [17] [18] [19] [20], mediating the switch from the DMN to a task-active state [18] [17].

In line with our previous findings in healthy controls that tDCS accentuates the brain network activity associated with the underlying brain state, we hypothesized that brain state is a core determinant of tDCS’s effects on brain networks in TBI patients. We further hypothesize that WM structure will additionally modulate the physiological effects of tDCS, with higher WM structural connectivity corresponding to greater brain network activity.

## Methods

### Participants

We recruited 59 participants, consisting of healthy controls (n=24; 12F:12M; mean age=39 years, SD=15.8 years) and TBI patients (n=35; 5F:30M; mean age=39.7 years, SD=10.3 years) (Table 1). Clinical characteristics of this cohort are previously described [11], and summarized in Table 1. Inclusion and exclusion criteria are described in (Table 2). One TBI participant was excluded due to distortion in fMRI preprocessing. Diffusion weighted MRI was collected from twenty-two healthy controls and all TBI patients. All participants were naïve to tDCS, and gave written informed consent. Ethical approval for the study was granted through the local ethics board (NRES Committee London-West London and GTAC).

**Table 1.**
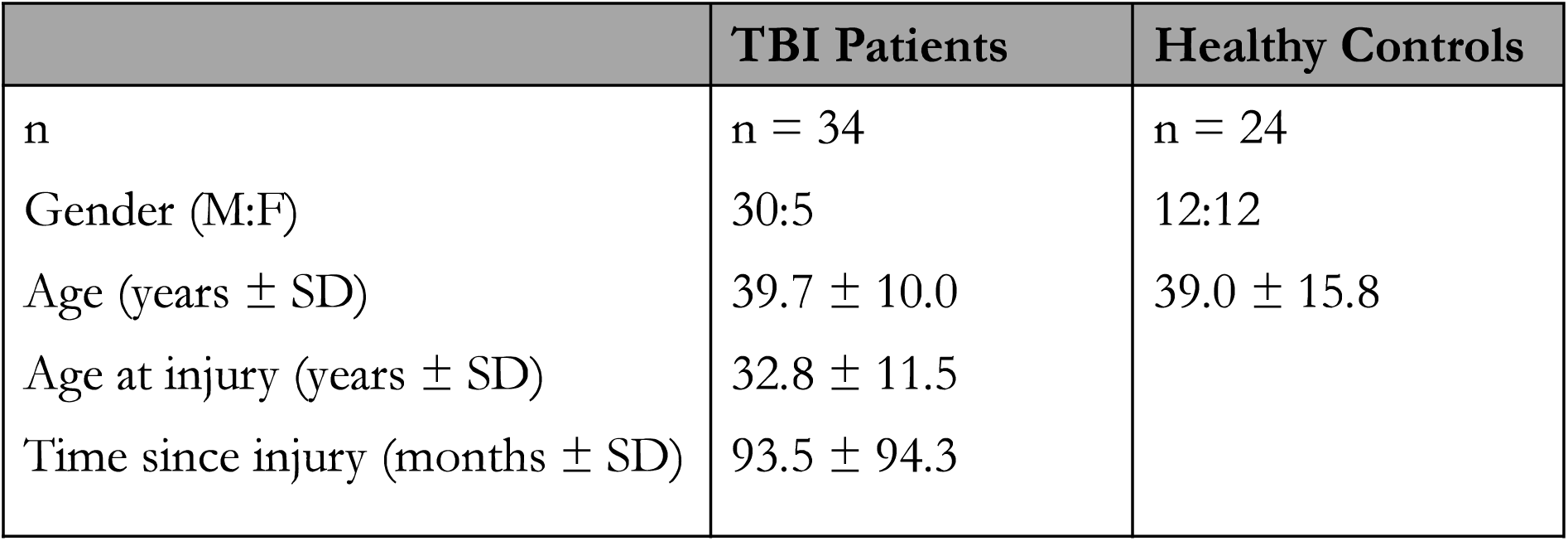
TBI patient and healthy control demographic information.

|  | TBI Patients | Healthy Controls |
| --- | --- | --- |
| n | n = 34 | n = 24 |
| Gender (M:F) | 30:5 | 12:12 |
| Age (years ± SD) | 39.7 ± 10.0 | 39.0 ± 15.8 |
| Age at injury (years ± SD) | 32.8 ± 11.5 | |
| Time since injury (months ± SD) | 93.5 ± 94.3 | |

**Table 2.** Participant inclusion and exclusion criteria

| Exclusion Criteria | Inclusion Criteria |
| --- | --- |
| Significant premorbid neurological or psychiatric illness, or learning disability<br>Substantial lesion directly affecting the right inferior frontal gyrus<br>Contraindication to tDCS (seizures within 12 months, broken skin over the site of stimulation, metallic implants in skull or eyes)<br>Cognitive impairment of such severity that the subject is unable to cooperate with the study<br>Contraindication to MRI scanning (including pregnancy) | A diagnosis of moderate-severe TBI (based on Mayo Clinic classification)<br>Able to provide informed consent for participation<br>At least 6 months since time of injury<br>Subjective or objective evidence of cognitive impairment after TBI<br>Clinically stable following TBI<br>Naïve to tDCS<br>Able to complete scanning protocol<br>Aged 16-80 years old |

### tDCS-fMRI paradigm

The Choice Reaction Task (CRT) consists of a random sequence of left or right arrows, to which participants had 1.3 seconds to respond by pressing a button with the index finger of the corresponding hand (Fig 1a). The tDCS-fMRI paradigm consisted of alternating blocks of CRT and “rest”. During the task and rest blocks participants received anodal, cathodal, or sham tDCS, creating six different conditions of combined brain state and stimulation polarity: “rest”+sham; “rest”+anodal; “rest”+cathodal; CRT+sham; CRT+anodal; CRT+cathodal (Fig 1b). Each run comprised 4 blocks of each condition, interspersed with brief black screen periods, presented in the same, pseudo-randomized order for all participants. Participants performed three runs interspersed with 2-3 minutes of rest to prevent fatigue. This resulted in a total of 18 minutes of full intensity tDCS. Prior to the tDCS-fMRI paradigm, participants performed a shorter version of the CRT. This was to confirm that the task produced the expected patterns of brain network activity in our population, and was used to create masks for the region of interest analysis evaluating the interaction between the brain state and polarity dependent on stimulation-induced brain network activity.

**Figure 1.**
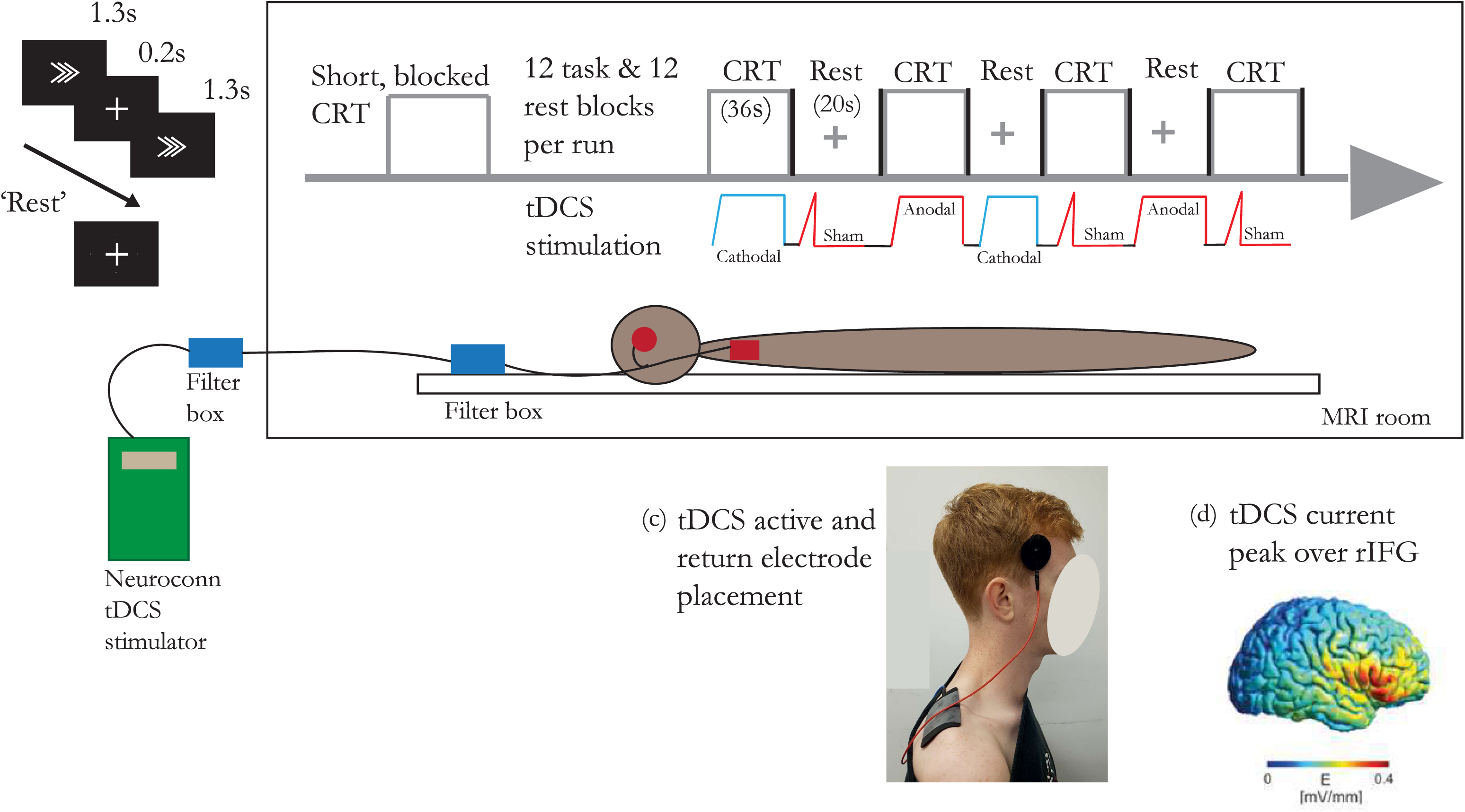
(a) Stimuli of the CRT and (b) experiment paradigm with concurrent tDCS/fMRI. Each participant performed three runs, each run consisting of four blocks of the six unique combined stimulation and task conditio ns (CRT + anodal, CRT+ cathodal, CRT+ sham, “rest” + anodal, “rest” + cathodal, “rest”+ sham). Blocks were interspersed by a blank screen with no stimulation. (c) Photo of the tDCS experimental setup. (d) Modelling of the electric field distribution over the rIFG.

### Delivery of tDCS

The active electrode was placed at F8 (10-20 EEG system) and the return electrode on the right shoulder (Fig 1c). Pre-stimulation impedances were below 3 k, and maximum impedance during stimulation was 29 k. To reduce impedance, electrodes were affixed with a layer of conductive paste (Ten20, D.O. Weaver, Aurora, CO). Anodal and cathodal stimulation were delivered using a 4.5 s ramp up, followed by full stimulation intensity at 1.8mA, then a 0.5 s ramp-down (Fig 1b). Sham stimulation consisted of ramp stages only. TTL triggers from the scanner were sent to the main computer, which controlled the simulation triggers via National Instruments DAQ device (National Instruments, Newbury, UK) using in-house MATLAB scripts. RF filtering boxes were located both inside the control room and the scanner bore and connected by Y-cable (RF-59/ULF; length=3 meters) and custom-made filter in the penetration panel to the stimulator in the control room. Wires from the participant and scanner were routed via the back of the bore to the control room. Tests used to ensure safe operating standards were conducted as per [21]. Peak electric field was confirmed via computational, finite element method (FEM) head modelling as in Li *et al* [9] (Fig 1d). Outside the scanner, participants had two 15-second blocks of anodal and cathodal tDCS, and were asked to rate sensations of itching, pain, metallic taste, burning, anxiety, and tingling on a 1-5 scale, with 1=nil and 5=unbearable. The average ratings for all categories found no difference between anodal or cathodal tDCS. Further details of the experimental paradigm and stimulation can be found in [9].

### Non-diffusion structural and functional MRI acquisition and preprocessing

T1 and fMRI sequences were acquired using a 3 T Siemens Verio (Siemens, Erlangen, Germany) and 32 channel head coil using parameters modeled from [21]. fMRI data preprocessing was performed as in [9] using FMRI Expert Analysis Tool (FEAT) Version 6.00, from FMRIB’s Software Library (FSL) [22]. Motion correction was performed using MCFLIRT [23]. FMRIB’s Nonlinear Image Registration Tool (FNIRT) was used to conduct removal of low-frequency drifts, spatial smoothing used a Gaussian kernel filter with a full width at half maximum of 6 mm, brain extraction to remove non-brain tissue (BET) [24], and coregistration. Using the participant’s T1-weighted scan and preprocessed data, participant’s fMRI volumes were registered to Montreal Neurological Institute (MNI) 152 standard space. Independent component analysis (ICA) analysis was conducted on each run using Multivariate Exploratory Linear Optimized Decomposition (MELODIC) [25]. Components were classified using a trained, automatic classification software, FMRIB’s ICA-based Xnoisefier (FIX) [26] and noise or non-brain components were removed.

### fMRI analysis: activation

A general linear model (GLM) was used to determine the relationship between activation and the task or “rest” conditions. In brief, single-session, subject level GLMs were first conducted to determine the effects of anodal and cathodal stimulation during the task, with the following regressors of interest: [all task block], [all anodal block], [all cathodal block]. The interactive regressors [task+anodal] and [task+cathodal] were run to demonstrate the interactive effects of stimulation polarity (anodal or cathodal) and brain state. The implicit baseline consisted of the [“rest”+sham] blocks. Subject levels were run again using [all rest block] rather than the [all task block] to demonstrate the effects of anodal and cathodal tDCS during the resting state. To interrogate the effects of stimulation in the absence of task a GLM was constructed substituting “rest” for task, resulting in [all “rest” blocks]. The interactive regressors [“rest”+anodal] and [“rest”+cathodal] then demonstrated the interactive effects of anodal and cathodal stimulation in the absence of task. All were run using square wave, double-gamma HRF in FSL’s FEAT. Six movement regressors were covaried out to account for motion artefact. A regressor of no interest to account for the periods of black screen was included. Group-level, mixed effects analysis combined all participant’s sessions, then all participants were combined using FLAME 1+2 in FSL FEAT. The parameter estimates [task+anodal], [task+cathodal], [“rest”+anodal], [“rest”+cathodal], [“rest”+anodal] >[“rest”+cathodal], and [task+anodal] >[task+cathodal] were run, as were the inverse estimates. Z statistical images were thresholded using Gaussian random field-based cluster inference with threshold of Z>3.1, corrected for cluster significance at a threshold of p=0.05. Further details can be found in [9] Determining the interaction between state and polarity dependent effects of tDCS on TBI patients was conducted with a region of interest (ROI) approach. The “task activated” and “task deactivated” network consisted of a binarized mask of the regions of increased (task activated) or decreased (task deactivated) BOLD activation during the shorter, non-stimulated, blocked CRT during the [task>“rest”] contrast.

### Diffusion Tensor Imaging (DTI) acquisition and analysis

Diffusion weighted imaging was collected from n=22 healthy control and n=33 TBI participants. Diffusion-weighted volumes were acquired using a 64-direction protocol (64 slices, in-plane resolution=2×2 mm, slice thickness=2 mm, field of view=25:6×25:6 cm, matrix size=128×128, TR=9500 ms, TE=103 ms, bvalue=0 mm^2^s^−1^). Four images were also acquired without diffusion weighting (b-value = 0 mm^2^s^−1^). DTI data were corrected for head motion and eddy current distortions, brain mask was extracted and constrained the tensor model using FSL’s FMRIB’s Diffusion Toolbox. Application of the tensor model generated voxelwise, individual FA maps. The maps were transformed into 1mm-resolution standard space using DTI-TK [27]. An initial group-based template was generated [28], and individual tensor-based images were then registered to the group template using diffeomorphic transformations. To assess WM structural connectivity in the whole brain and of regions composing the two brain networks of interest (SN and DMN) FA values were extracted for the following structures as obtained in [29]:

1. The Whole Skeleton tract assesses whole-brain WM tract integrity.
2. The rAI-dACC/pre-SMA tract assesses SN structural integrity in the tract connecting the right anterior insula (rAI) to the dorsal anterior cingulate (dACC) / pre-supplementary motor area (pre-SMA). This tract partly overlaps the frontal Aslant tract described by [30].
3. The mPFC-PCC/PRE (medial prefrontal cortex to posterior cingulate cortex/precuneus) tract assessed DMN structural integrity within the bilateral cingulum.

The influence of white matter structure on the brain state and polarity dependent effects of stimulation were determined using FA as a continuous covariate, in higher-level FEAT analyses as single-group averages with continuous covariate interaction. A ROI approach was also used to determine the relationship between the physiological effects of stimulation and WM structure. The areas of activity influenced by WM structure were used to create a binarized mask. fslstats were used to extract β values from the mask for the areas of activity. A Spearman correlation was used to determine the relationship between the β values and FA, and was Bonferroni corrected for multiple comparisons.

### Statistical analysis of behavioral results

Statistical analysis was conducted using MATLAB [31] and R (www.r-project.org). Kolgov-Smirnov tests evaluated normalcy of data.

## Results

### The brain network effects of TDCS are dependent on cognitive brain state in both healthy and TBI participants

In both healthy and TBI participants, CRT performance without stimulation displayed robust activation across fronto-parietal regions, including the dACC node of the SN, primary sensory/motor cortex, basal ganglia, and bilateral thalami (Fig 2a). There was concurrent deactivation of the posterior cingulate cortex (PCC) node of the DMN.

**Figure 2.**
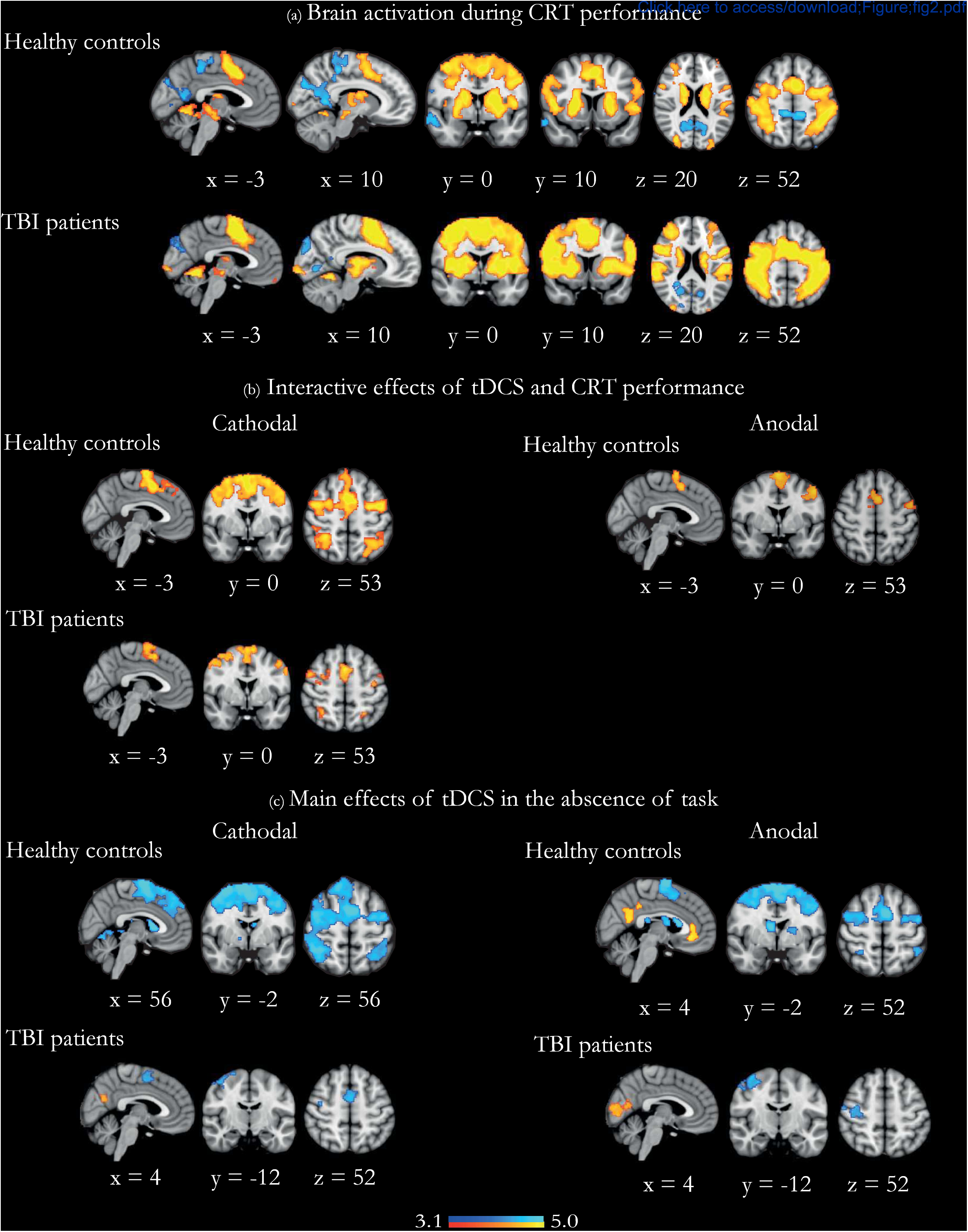
The physiological effects of tDCS during and in the absence of the Choice Reaction Task. (a) Overlay of brain activation (in warm colors) and deactivation (in cool colors) during the short CRT with no tDCS, in healthy and TBI participants. (b) Brain regions with further activation due to anodal and cathodal stimulation applied during CRT performance in healthy participants, and brain regions with further activation due to cathodal stimulation applied during CRT performance in TBI participants. (c) Brain regions showing the main effect of anodal and cathodal stimulation in the absence of task in healthy and TBI participants. Warm colors denote further activation, and cool colors denote further deactivation. Results are superimposed on a Imm MNI152 template. Cluster corrected z = 3.1, p < 0.05.

The interactive effects of tDCS and CRT performance and the main effect of stimulation in the absences of task have been previously described in healthy participants [9]. In essence, both anodal and cathodal stimulation act to accentuate the underlying patterns of brain network activity set by the current cognitive brain state. That is, the interactive effect of anodal or cathodal tDCS and CRT performance was additional increased activation in the dACC/pre-SMA and lateral prefrontal regions, which are all areas already activated during CRT performance (Fig 2b). Conversely, the main effect of anodal and cathodal tDCS in the absence of task was increased deactivation of the rIFG and underlying anterior insula, bilateral superior frontal gyri, bilateral frontal eye fields, and bilateral superior parietal regions, areas which are already active in the absence of task. An additional main effect of anodal tDCS in the absence of task was further activation of the PCC, the precuneus, and the ventromedial prefrontal cortex (vmPFC), areas associated with the DMN (Fig 2c).

In TBI participants, the interactive effect of cathodal tDCS and CRT performance was additional increased activation in the dACC node of the SN and, to a lesser extent, primary sensory/ motor cortex, basal ganglia, and bilateral thalami, which are all areas already activated during CRT performance (Fig 2b). There was no interactive effect of anodal tDCS and CRT performance over and above the brain activity patterns elicited by CRT performance itself. Conversely, the main effect of cathodal tDCS in the absence of task was increased activation in PCC and further deactivation of dACC, and the main effect of anodal tDCS in the absence of task was increased activation in medial occipital cortex and further deactivation of sensory/motor cortex (Fig 2c).

These findings in the TBI cohort almost entirely replicate our previous work in healthy participants [9]. Indeed, there were no whole-brain differences when contrasting TBI and healthy participants in any of the five conditions presented in Fig 2.

### The brain state-dependent effects of tDCS in TBI patients are different between anodal and cathodal stimulation

An ROI approach was used to investigate the interaction between the brain state and polarity dependent effects of tDCS [9], focusing analysis to brain network areas relevant to CRT performance. We found a significant interaction between brain state and stimulation polarity in both the task-activated regions (F [1,213]=4.23, p=0.042) (Fig 3a) and task-deactivated regions (F [1,412]=4.16, p=0.043) (Fig 3b).

**Figure 3.**
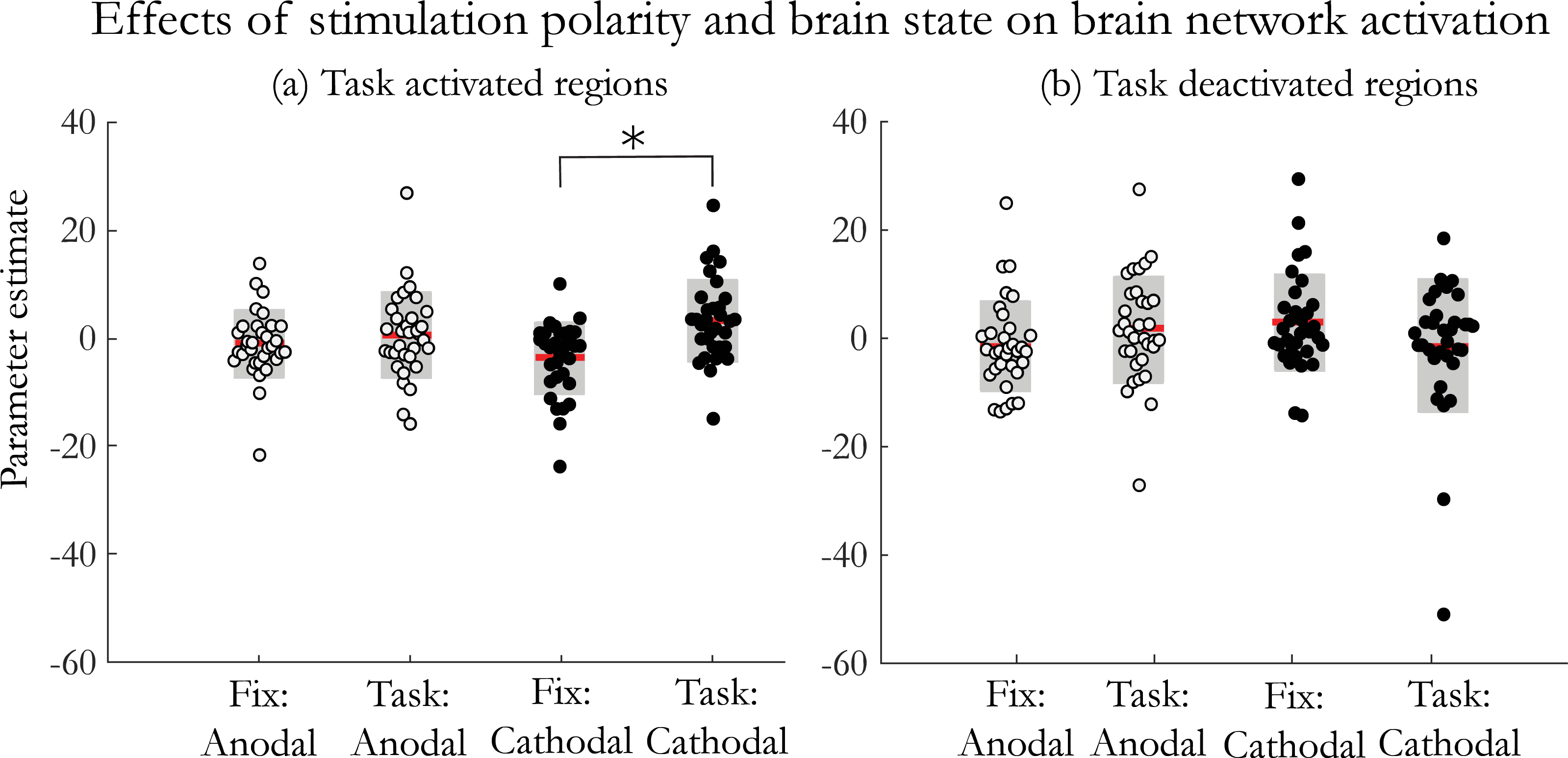
The interaction between brain state and stimulation polarity on network activation. Charts showing changes in BOLD activit y in task-related networks under anodal and cathodal tDCS during CRT performance (labeled ‘Task’) and in the absence of task (labeled ‘Fix’). (a) “Task activated regions” refers to brain regions showing increased activation, and (b) “Task deactivated regions” refers to brain regions showing decreased activation during CRT performance, as shown in Figure 2a. White dots indicate anodal stimulation, black dots are for cathodal. Red lines are group mean values. Grey boxes are ± 1 standard deviation from the mean. * denotes p < 0.05.

During both task and fixation, both stimulation polarities modulated BOLD activity to emphasize underlying network activity. Post-hoc t-tests confirmed cathodal stimulation during task produced greater BOLD activity than activation due to cathodal stimulation in the absence of task (t(11.84)=2.384, p=0.004) (Fig 3b). Similarly, cathodal stimulation during task produced greater BOLD activity than activation in the absence of task due to anodal stimulation (t(8.65)=0.782, p=0.020). In the absence of task, activity due to cathodal stimulation was less pronounced than anodal stimulation during task (t(8.56)=0.448, p=0.031). There was no difference between the effects of anodal and cathodal stimulation in the task-activated regions. Overall, the marked effects seen in cathodal stimulation were not seen with anodal stimulation.

### WM structure further modulates the brain state-dependent effects of tDCS on brain network activity

TBI participants had reduced mean FA within the whole skeleton [t(52)=2.299, p=0.0256] in the rAI-dACC/pre-SMA tract representing the SN [t(52)=3.137, p=0.0028] and in the cingulum bundle, representing the DMN [t(32)=2.535, p=0.0143] (Fig 4). As there were no group differences in the brain-state dependent effects of tDCS, TBI patients and healthy controls were combined to determine the influence of WM structure on the physiological effects of stimulation (Fig 5a).

**Figure 4.**
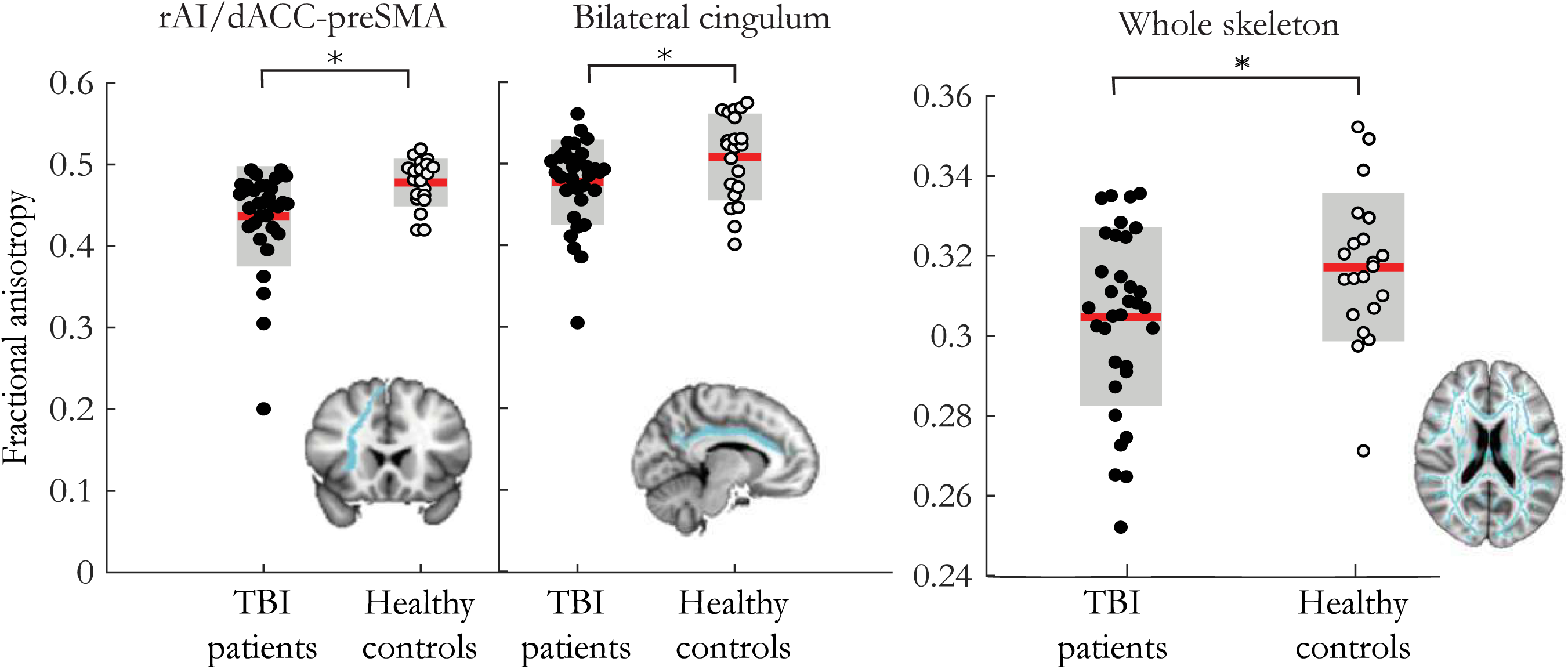
WM structure in key tracts in TBI patients and healthy controls. Red lines are group mean values. Grey boxes are ± 1 standard deviation from the mean. Inset brain pictures show rAI-dACC/pre-SMA, bilateral cingulum and whole skeleton tracts. * denotes p < 0.05.

**Figure 5.**
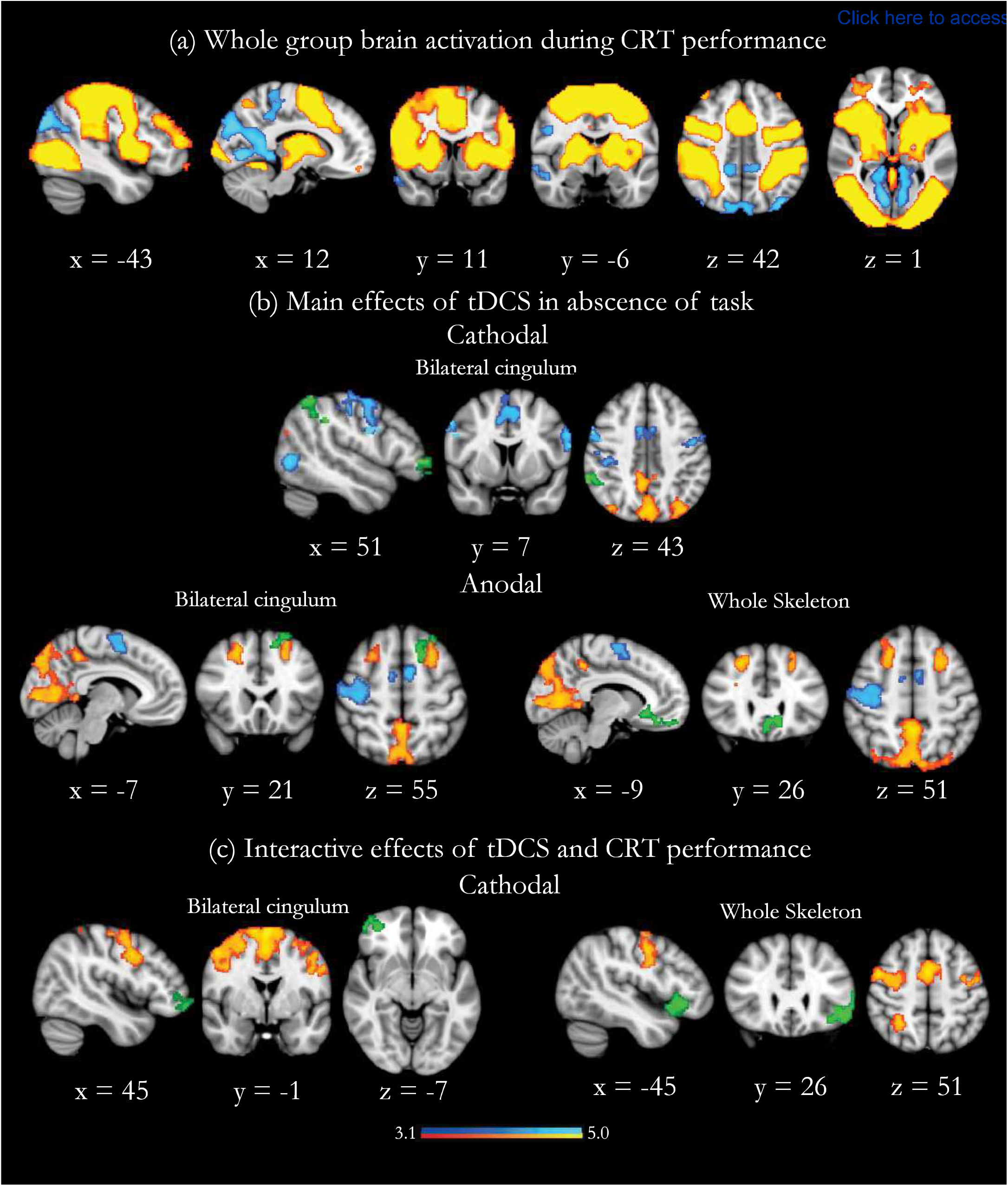
The influence of WM structure on physiological effects of tDCS during and in the absence of task. (a) Overlay of brain activation (in warm colors) and deactivation (in cool colors) during CRT performance in the whole group (TBI patients and healthy controls. b) Brain regions where WM structure further modulates stimulation-induced BOLD activity in the absence of task (in green). Warm and cool colors denote the BOLD activity produced by stimulation, without accounting for white matter structure. (c) Brain regions where WM structure further modulates stimulation-induced BOLD activity during task performance (in green). Warm and cool colors denote the BOLD activity produced by stimulation, without accounting for white matter structure. Results are superimposed on a Imm MNI152 template. Cluster corrected z = 3.1, p < 0.05.

#### In the Absence of Task

Bilateral cingulum FA influenced BOLD response within several regions to cathodal stimulation: the right frontal pole, middle frontal gyrus, and inferior parietal lobule (Fig 5b). Increases in bilateral cingulum FA correlated with increases in activity in these regions (r_s_=0.446, p=8.25 e-04) (Fig 6a). Whole skeleton FA modulated BOLD response within the vmPFC during anodal stimulation in the absence of task (Fig 5b). As whole skeleton FA increases, vmPFC activation increases (r_s_=0.500, p=1.44 e-04) (Fig 6a). Bilateral cingulum FA influenced BOLD response within the left middle frontal gyri to anodal stimulation in the absence of task (Fig 5b). As bilateral cingulum FA increases, so does activation in the left middle frontal gyri (r_s_=0.361, p=0.008) (Fig 6a). In summary, there were positive correlations between WM structure and brain network activation in the absence of task. There was no influence of whole skeleton FA on brain activity due to cathodal stimulation, and there was no influence of rAI-dACC/preSMA FA on brain activity due to anodal or cathodal stimulation.

**Figure 6.**
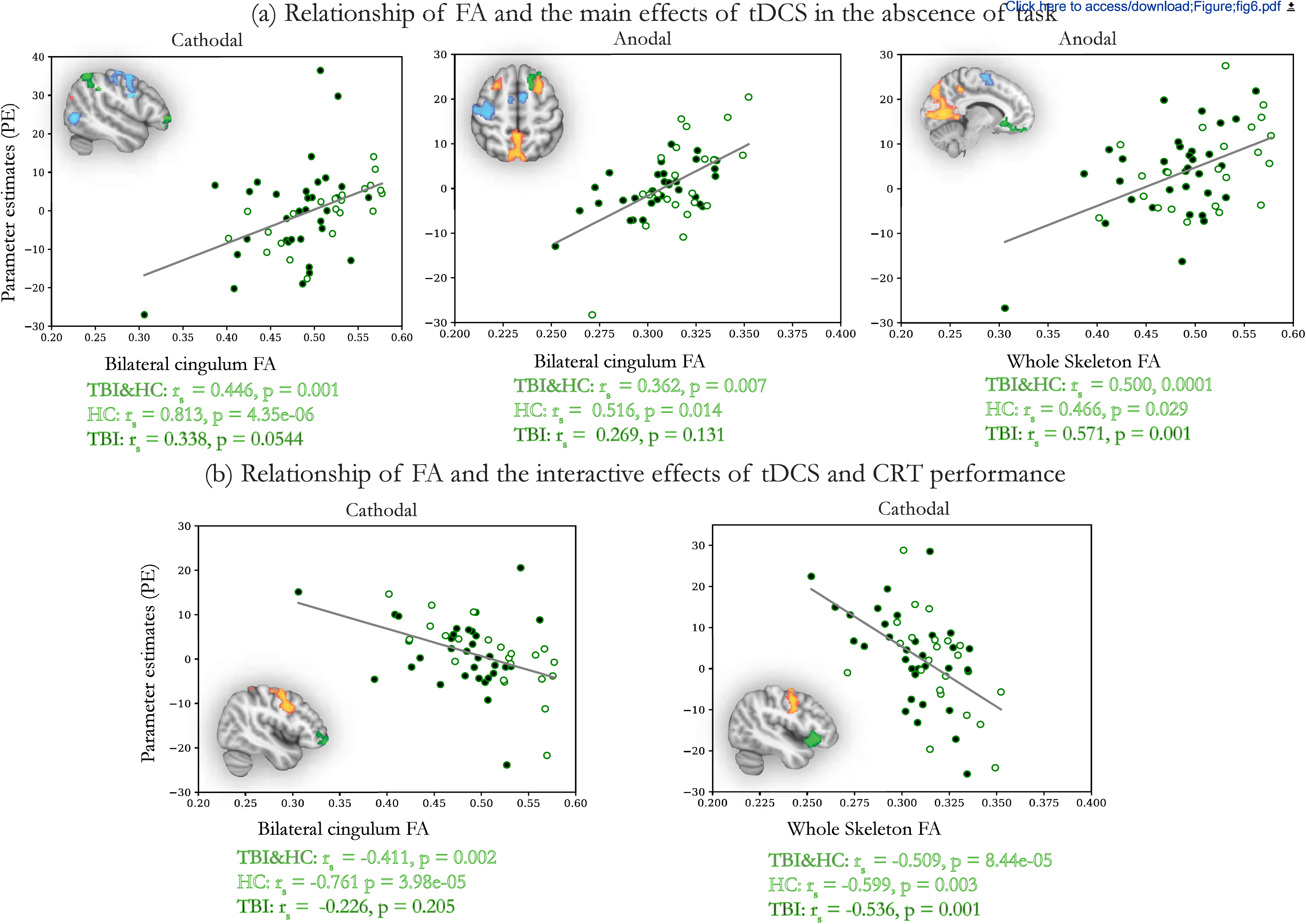
The influence of WM structure on physiological effects of tDCS during and in the absence of task. Scatter plots demonstrate the nature of the relationship between WM structure and stimulation-induced BOLD activity. (a) Relationship between FA and the main effect of tDCS in the absence of task for TBI patients (black circles) and healthy controls (white circles). BOLD activity was extracted from the green regions in the brain insets, which correspond to the results shown in Fig 5b. (b) Relationship between FA and the interactive effects of cathodal tDCS and CRT performance in TBI for TBI patients (black circles) and healthy controls (white circles). BOLD activity was extracted from the green regions in the brain insets, which correspond to the results shown in Fig 5c. For all plots, FA is on the x-axis, and parameter estimates indicate the level of BOLD activity on the y-axis. Line of best fit is fitted to the whole-group data. Spearman correlations for the whole-group, and each group separately, are shown below the plots.

**Figure 7.**
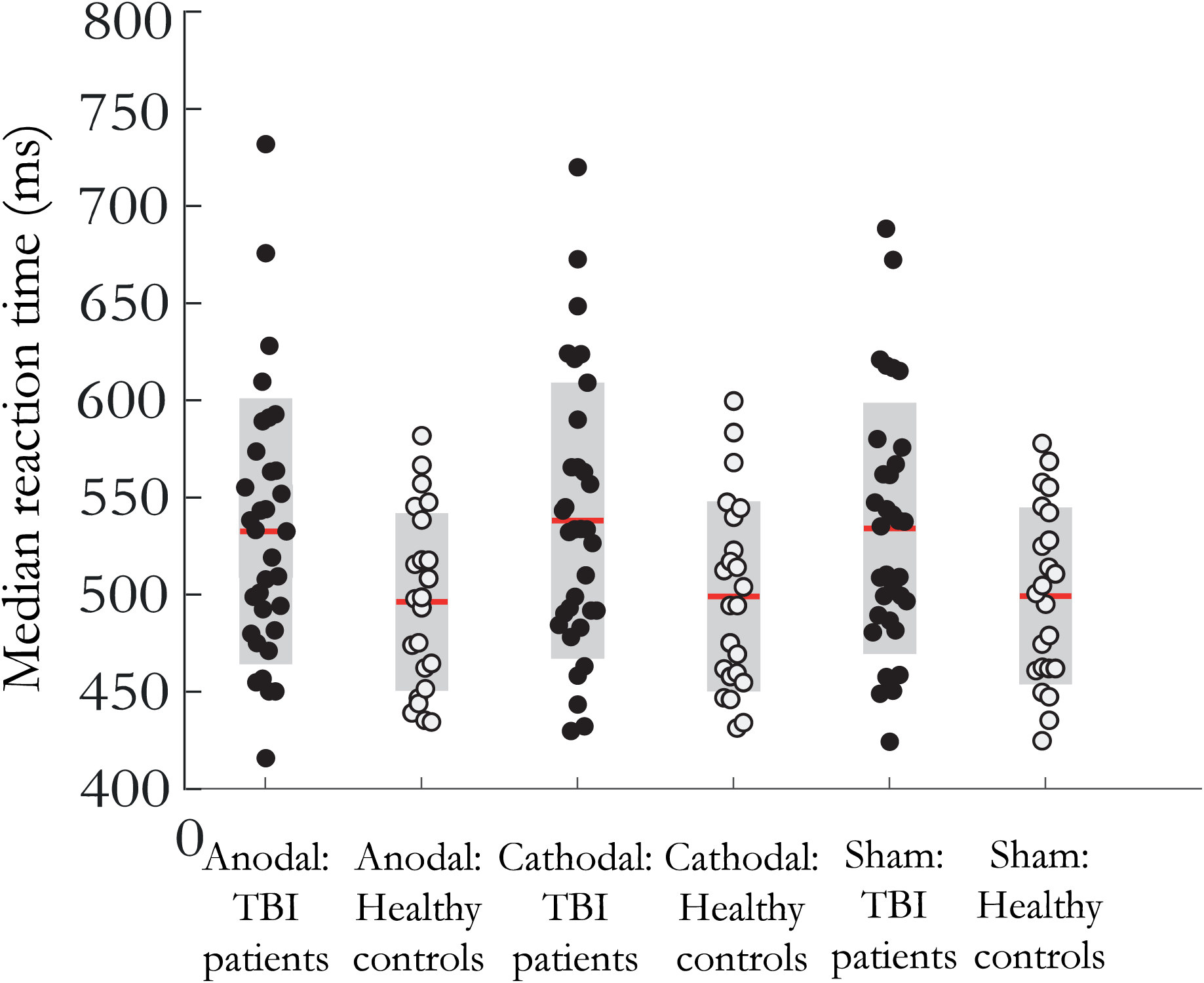
Median reaction time on the CRT in healthy controls (open circles) and TBI patients (filled circles) with anodal, cathodal, and sham stimulation. Red lines are group mean values. Grey boxes are ± 1 standard deviation from the mean.

#### During Task Performance

Whole skeleton FA influenced the BOLD response within the left IFG with cathodal stimulation applied during task performance (Fig 5c). As whole skeleton FA increased, there was more left IFG deactivation (r_s_=0.433, p=0.001) (Fig 6b). Bilateral cingulum FA influenced the BOLD response within the right IFG/ right frontal pole with cathodal stimulation applied during task. As bilateral cingulum FA increases, there was more right IFG/frontal pole deactivation (r_s_=0.433, p=0.001) (Fig 6b). In summary, there was an inverse relationship between WM structure and brain network activity with tDCS applied during task performance. There was no influence of rAI-dACC/preSMA FA on brain activity during cathodal stimulation. There was no influence of FA from any of the tracts on brain activity during anodal stimulation.

### Behavioral performance in the Choice Reaction Task

There was a main effect of group on median reaction time (medRT) (F[1,13.57]=168, p=0.0003), with TBI participants having a longer medRT than healthy participants. There was no main effect of group on accuracy (F[1,2.556]=171, p=0.112), with both groups performing at >90% accuracy in all conditions (Fig 7). There was no main effect of stimulation type on medRT or accuracy, and no interaction between group and stimulation type on medRT or accuracy (all p>0.05).

## Discussion

We show that brain state is a key factor that determines the physiological effects of tDCS in TBI participants. This replicates findings from our previous study in a separate, clinical cohort. We further show that WM structure modulates the influence of brain state on brain network effects of tDCS. Our results show how participant and experimental factors interact to shape the physiological effects of tDCS, and extend our mechanistic understanding of tDCS’s effect on brain network dynamics.

### Replicating the brain state dependent effects of tDCS

We have previously shown that brain network effects of tDCS are highly dependent on underlying cognitive brain state [9]. We now replicate this finding in a completely new, patient cohort, which suggests that the brain-state dependence of tDCS action is a core characteristic of tDCS-induced neural activity. As in healthy controls, tDCS seems to reinforce underlying brain network activity. During the CRT, cathodal stimulation increases SN activity over and above normal SN activity during the task. In the absence of task, the same stimulation polarity actually produced decreased SN activity, thus reinforcing state-dependent patterns of brain network activity. We have shown the state-dependent effects of tDCS in both healthy control [9] and now a patient population. This suggests that brain state is a particularly important determinant of tDCS’s effects and is broadly generalizable across populations. This framework seems to generalize to other modalities [32] [33]. For example, a study studying tDCS-induced changes in cortical excitability, as assessed by motor evoked potential, found the physiological effects of tDCS were dependent on brain state [33]. This diverges from the anodal-excitatory, cathodal-inhibitory dichotomy seen on a cellular level [34]. It has been hypothesized that the different effects on the macro/meso level compared to the microscopic level could result from tDCS’s effects on a cellular level operating on various excitatory and inhibitory populations, shifting the balance of large-scale, state-dependent brain networks [35].

#### The brain state-dependent effects of tDCS on brain network activity interact with WM structure

We found that WM structure influences tDCS-induced brain activity in a state-dependent manner. In the absence of task, higher FA was correlated with increased brain activity in vmPFC, rIFG, middle frontal gyrus, and IPL. These areas, particularly IPL and vmPFC, are part of the DMN, and usually show increased activity in the absence of CRT performance for both anodal and cathodal stimulation. We have already shown that the effects of tDCS are brain state dependent, and now find that, in the absence of task, WM structure seems to accentuate the state-dependent response to tDCS.

One possible reason for the state-dependent directionality of WM structure’s influence on brain network activity might be the extent of structural connectivity to the node itself, and its functional role in varying brain network dynamics. In the absence of task, we found bilateral cingulum FA influenced increased activity in the IPL, left middle and frontal gyri. The IPL is a component of the DMN and FPCN, and has been identified for its role in attentional control [36]. This explains why bilateral cingulum FA influences increased activation of the IPL-a region with high functional and structural overlap with the DMN and SN. A recent computational simulation study investigated the influence of structural and functional connectivity of a node’s ability to change brain network activity upon stimulation [37]. They found that stimulating the IPL drives brain network activity to a target state, due partially to its high structural connectivity [37] [38]. This strengthens the theory that in the absence of task WM structure influences the brain state-dependent physiological effects of tDCS.

Conversely, during task performance, there was a negative relationship between FA and brain network activity, seen in bilateral IFGs. These are regions which, in our population, show an increase in activity during CRT performance. We have previously shown that healthy controls in a high FA group demonstrated greater rIFG and right anterior insula activation than a low FA group during anodal tDCS during a different, more complex cognitive task [10]. This led us to predict that higher FA would result in increased rIFG/left IFG activation with tDCS during CRT performance. This unexpected finding suggests that during CRT performance the relationship between WM structure and tDCS-induced brain activity is no longer injective. One possible extension of this study would be to vary the degree of task difficulty as a way to continuously manipulate brain state, to map out the interaction between WM structure and brain state in more detail.

Therefore, we see that the same stimulation polarity (cathodal) and same WM tract (bilateral cingulum) seem to shape the brain network effects of stimulation in different ways depending on brain state. This leads us to speculate that there is a ‘hierarchy of influences’ on brain network activity response to tDCS, where some factors are more influential than others in determining brain network response to tDCS.

#### Limitations

Our stimulation target was limited to the rIFG. Though we chose the rIFG for its unique role in the SN/DMN dynamics, we postulate stimulation of other nodes in the DMN or SN- such as the IPL-would encourage similar network driven results. The pseudorandomized nature of our study was a precaution against carry-over effects of stimulation. If there is some carryover, the pseudorandomisation is more likely to produce noise rather than systematic bias.

#### Conclusion

We provide further evidence for the important influence of brain state on the brain network effect of tDCS. We further show that WM structure interacts with brain state in its effect on tDCS-induced brain network activity. It is possible that participant and experimental factors follow a ‘hierarchy of influence’, with factors such as brain state exerting a greater, more generalizable influence on the physiological effects of tDCS because it is higher up on the ‘hierarchy’. Further studies specifically investigating parameter interactions on tDCS-induced brain network activity will help improve our ability to translate the use of stimulation.

## Funding

This work had the following funding: LML was funded by Wellcome Trust Clinical Research Training Fellowship (103429/Z/13/Z) and the NIHR Academic Clinical Lectureship program. I.R.V. was funded by the Wellcome Trust (Ref: 103045/Z/13/Z) and the BBSRC (Ref: BB/S008314/1), DJS is funded by a National Institute for Health Research (NIHR) Professorship.

## Acknowledgements

We would like to thank all participants. We would like to thank Dr Jonathan Howard for tireless and insightful technical and methodological assistance, and Dr Alexander Optiz for doing the modelling for peak electric field. This study was conducted at the Imperial College Clinical Imaging Facility. The study was supported by the NIHR Imperial College London Biomedical Research Centre and the Centre for Blast Injury Studies (CBIS) at Imperial College London.

## Author Approval

All authors have seen and approved this manuscript. There are no competing interests to disclose. As of the date of submission to BioRix, this manuscript has not been published nor accepted elsewhere.

## Abbreviations

SN: salience network
FPCN: frontoparietal control network
DMN: default mode network
rIFG: right inferior frontal gyrus
dACC/pre-SMA: dorsal anterior cingulate/pre-supplementary motor area
PCC: posterior cingulate cortex
vmPFC: ventromedial prefrontal cortex

## Notes

### Competing Interest Statement

The authors have declared no competing interest.

